## Supplemental Table 1 for "Long-term alterations in brain and behavior after postnatal Zika virus infections in infant macaques" (from another version of this preprint: version 1, 2019-10-01)

**Supplemental Table 1.**
**Social Behavior with Familiar Partner in Novel Play Cage**

| Behavior | Subject | Day 1 | Day 2 | Day 3 | Day 4 | Day 5 | Day 6 |
|---|---|---|---|---|---|---|---|
| Latency to Enter Play Cage (seconds) | Control-1 | 10.68 | 1.50 | 14.45 | 0.70 | 1.23 | 0.87 |
| | Control-2 | 29.80 | 18.05 | 1.50 | 9.54 | 0.97 | 5.07 |
| | ZIKV-5 | 169.14 | 143.97 | 139.40 | 29.70 | 2.70 | 5.51 |
| | ZIKV-6 | 105.14 | 208.74 | 147.91 | 40.24 | 10.71 | 7.54 |
| Percent Time Spent Together with the Familiar Partner | Control-1 | 93.27 | 93.24 | 86.19 | 86.89 | 86.87 | 87.70 |
| | Control-2 | 93.27 | 93.24 | 86.19 | 86.89 | 86.87 | 87.70 |
| | ZIKV-5 | 90.92 | 66.37 | 64.90 | 61.57 | 66.49 | 40.62 |
| | ZIKV-6 | 90.92 | 66.37 | 64.90 | 61.57 | 66.49 | 40.62 |
| Percent Time Spent Away from the Familiar Partner | Control-1 | 6.73 | 6.76 | 13.82 | 13.11 | 13.14 | 12.31 |
| | Control-2 | 6.73 | 6.76 | 13.82 | 13.11 | 13.14 | 12.31 |
| | ZIKV-5 | 9.08 | 33.63 | 34.92 | 38.43 | 33.52 | 59.38 |
| | ZIKV-6 | 9.08 | 33.63 | 34.92 | 38.43 | 33.52 | 59.38 |
| Percent Time Spent Along the Perimeter of the Play Cage | Control-1 | 18.89 | 28.25 | 50.06 | 65.51 | 71.64 | 72.17 |
| | Control-2 | 4.76 | 16.99 | 34.37 | 46.64 | 51.27 | 43.58 |
| | ZIKV-5 | 97.47 | 95.65 | 96.48 | 94.55 | 96.95 | 90.10 |
| | ZIKV-6 | 96.10 | 98.52 | 99.22 | 93.57 | 98.08 | 94.02 |
| Percent Time Spent in Center of the Play Cage | Control-1 | 80.45 | 71.12 | 46.84 | 34.55 | 28.41 | 28.04 |
| | Control-2 | 95.24 | 82.96 | 65.62 | 53.70 | 48.77 | 56.61 |
| | ZIKV-5 | 2.49 | 4.28 | 3.53 | 5.45 | 3.15 | 10.12 |
| | ZIKV-6 | 1.67 | 1.48 | 0.78 | 6.43 | 2.30 | 6.28 |
| Loss of Balance (rate/min) | Control-1 | 0.085 | 0.084 | 0.000 | 0.043 | 0.108 | 0.130 |
| | Control-2 | 0.021 | 0.000 | 0.127 | 0.043 | 0.043 | 0.086 |
| | ZIKV-5 | 0.063 | 0.125 | 0.305 | 0.318 | 0.409 | 0.363 |
| | ZIKV-6 | 0.124 | 0.319 | 0.692 | 0.404 | 0.345 | 0.411 |
| Play Behavior (rate/min) | Control-1 | 0.076 | 0.070 | 0.058 | 0.07 | 0.086 | 0.096 |
| | Control-2 | 0.019 | 0.044 | 0.089 | 0.04 | 0.065 | 0.064 |
| | ZIKV-5 | 0.027 | 0.070 | 0.023 | 0.02 | 0.171 | 0.150 |
| | ZIKV-6 | 0.033 | 0.078 | 0.034 | 0.01 | 0.065 | 0.078 |
| Affiliative Behavior (rate/min) | Control-1 | 0.296 | 0.399 | 0.530 | 0.494 | 0.237 | 0.194 |
| | Control-2 | 0.806 | 0.443 | 0.233 | 0.215 | 0.237 | 0.346 |
| | ZIKV-5 | 0.169 | 0.000 | 0.000 | 0.000 | 0.000 | 0.000 |
| | ZIKV-6 | 0.435 | 0.106 | 0.041 | 0.085 | 0.000 | 0.048 |
| Hostile Behavior (rate/min) | Control-1 | 0.190 | 0.168 | 0.297 | 0.451 | 0.538 | 0.734 |
| | Control-2 | 0.403 | 0.274 | 0.211 | 0.172 | 0.409 | 0.324 |
| | ZIKV-5 | 0.381 | 0.852 | 1.097 | 2.080 | 1.785 | 2.516 |
| | ZIKV-6 | 0.331 | 1.084 | 0.610 | 1.021 | 0.711 | 0.823 |

| | | | | | | | |
|---|---|---|---|---|---|---|---|
| Submissive Behavior (rate/min) | Control-1 | 0.000 | 0.042 | 0.000 | 0.021 | 0.000 | 0.022 |
| | Control-2 | 0.000 | 0.148 | 0.021 | 0.000 | 0.000 | 0.022 |
| | ZIKV-5 | 0.000 | 0.062 | 0.081 | 0.149 | 0.215 | 0.194 |
| | ZIKV-6 | 0.083 | 0.191 | 0.102 | 0.064 | 0.172 | 1.016 |
| Anxious Behavior (rate/min) | Control 1 | 0.296 | 1.345 | 0.975 | 1.117 | 2.282 | 1.231 |
| | Control 2 | 0.021 | 0.380 | 0.106 | 0.172 | 0.560 | 0.302 |
| | ZIKV 5 | 0.042 | 0.083 | 0.020 | 0.085 | 0.000 | 0.000 |
| | ZIKV 6 | 0.041 | 0.043 | 0.020 | 0.043 | 0.043 | 0.145 |
| Vocalizations (rate/min) | Control 1 | 1.648 | 1.744 | 1.166 | 0.988 | 0.495 | 1.641 |
| | Control 2 | 0.085 | 0.084 | 0.021 | 0.129 | 0.431 | 0.713 |
| | ZIKV 5 | 0.909 | 0.104 | 0.183 | 0.021 | 0.108 | 0.266 |
| | ZIKV 6 | 0.455 | 0.043 | 0.000 | 0.043 | 0.065 | 0.436 |
| Stereotypies (rate/min) | Control 1 | 0.000 | 0.000 | 0.021 | 0.000 | 0.000 | 0.000 |
| | Control 2 | 0.713 | 0.764 | 0.823 | 0.151 | 0.538 | 0.302 |
| | ZIKV 5 | 0.266 | 0.021 | 0.000 | 0.000 | 0.000 | 0.000 |
| | ZIKV 6 | 0.103 | 0.000 | 0.000 | 0.000 | 0.000 | 0.000 |

List of behaviors coding during social interaction with a highly familiar partner in a large play cage and the individual values for each subject across the 6 days of testing.
