## Supplemental Table 4 for "Long-term alterations in brain and behavior after postnatal Zika virus infections in infant macaques" (from another version of this preprint: version 1, 2019-10-01)

**Supplemental Table 4. Behavioral Ethogram**

| Category and Specific Behaviors | Measurement | Brief Descriptions |
|---|---|---|
| Latency to Enter | Duration | Time between the door opening and the animal entering into the social play cage |
| Location in the Cage | | |
| Perimeter | Duration | Time spent along the walls and outside area of the social play cage |
| Center | Duration | Time spent on the floor, ceiling, or center pole of the social play cage |
| Motor Related Behaviors | | |
| Loss of Balance | Frequency | Uncoordinated movement that resulted in a brief loss of balance or footing |
| Play Behavior | Cumulative Duration | |
| Exploration | Duration | Use of hands or mouth to nonaggressively grab, hold, or inspect an object, toy, or social play cage |
| Swing/Jump | Duration | Vigorously jumping or swinging on objects, toys, or part of the social play cage |
| Distance to Partner | | |
| Together | Duration | Time animals spent in full body contact, proximity (arms length), or within 5 feet of each other |
| Away | Duration | Time animals spent more than 5 feet apart from each other |
| Affiliative | Cumulative Frequency | |
| Touch | Frequency | One animal uses its hand to briefly contact or hold onto the other animal (nonaggressively) |
| Groom | Frequency | One animal uses its hands or mouth to comb through the fur of the other animal |
| Follow | Frequency | One animal persistently trails the other, both are actively in motion |
| Aggression | Cumulative Frequency | |
| Threat bark | Frequency | Low pitch, high intensity, rasping, guttural |
| Threat - noncontact | Frequency | Any of the following: open mouth (no teeth exposed), head-bobbing, or ear flapping |
| Threat - contact | Frequency | Any of the following: bitting, slapping, hitting, or grabbing the other animal |
| Cage aggression | Frequency | Vigorously slaps, shakes, or slams body against cage |
| Chase | Frequency | Vigorous and rapid pursuit of the other animal |
| Submissive | Cumulative Frequency | |
| Withdrawal | Frequency | Quick, jerky motion away from the stimulus object (jump back) |
| Present | Frequency | Rigid posture with rump and tail elevated and oriented toward the other animal |
| Grimace[1] | Frequency | Refracted lips, exposed clenched teeth |

| | | |
|---|---|---|
| Anxiety | Cumulative Frequency | |
| Scratch | Frequency | Rapid scratch of body with hands or feet |
| Body shake | Frequency | Shake of the whole body or just head and shoulders region |
| Tooth grind[1] | Frequency | Repetitive, audible rubbing of upper and lower teeth |
| Yawn[1] | Frequency | Open mouth widely, exposing teeth |
| Vocalizations | Cumulative Frequency | |
| Coo | Frequency | Clear soft pitch and intensity, sounds like “ooooh” |
| Scream | Frequency | High-pitched vocalization, with extreme high intensity; sounds like “eeeeeee” |
| Stereotypies | Cumulative Duration | |
| Pacing | Duration | Repetitive motor pattern around the test cage |
| Motor stereotypy | Duration | Repetitive, abnormal voluntary or involuntary motor patterns (e.g. swinging, twirling, flipping) |

List of all behaviors scored, how they are measured and a brief definition. 1. Behavior that was not exhibited by any animal during observations.
