## Supplemental Table 3 for "Long-term alterations in brain and behavior after postnatal Zika virus infections in infant macaques" (from another version of this preprint: version 1, 2019-10-01)

### Supplemental Table 3.
### Functional Connectivity (r-values) Between Brain Regions of Interest

| | Subject | | | |
|---|---|---|---|---|
| **Regions of Interest** | **Control-1** | **Control-2** | **ZIKV-5** | **ZIKV-6** |
| amygdala-amygdala 12mo | 0.40446 | 0.26000 | 0.20683 | 0.34853 |
| RT amygdala-hippocampus 12mo | 0.37978 | -0.05650 | -0.06897 | 0.43040 |
| LT amygdala-hippocampus 12mo | 0.42025 | 0.55852 | 0.15477 | 0.28407 |
| RT amygdala-PFC32 12mo | -0.07096 | 0.12485 | -0.11577 | 0.13632 |
| LT amygdala-PFC32 12mo | 0.00266 | -0.13709 | 0.07632 | 0.03439 |
| RT amygdala-PFC46 12mo | 0.00140 | 0.02658 | 0.11837 | 0.11025 |
| LT amygdala-PFC46 12mo | -0.02930 | -0.08735 | 0.22407 | 0.03536 |
| RT amygdala-OFC11&13 12mo | -0.11172 | 0.05670 | -0.15514 | -0.12953 |
| LT amygdala-OFC11&13 12mo | 0.11385 | -0.04219 | 0.10252 | 0.08146 |
| RT amygdala-caudate 12mo | -0.00858 | 0.00351 | 0.11016 | 0.18491 |
| LT amygdala-caudate 12mo | -0.01131 | -0.00591 | 0.19473 | 0.06599 |
| RT hippocampus-OFC14 12mo | 0.10883 | 0.02029 | -0.15121 | 0.01819 |
| LT hippocampus-OFC14 12mo | 0.14742 | 0.04789 | 0.00319 | 0.06395 |
| RT hippocampus-PFC24 12mo | -0.10220 | 0.09304 | 0.11650 | 0.23022 |
| LT hippocampus-PFC24 12mo | 0.04504 | 0.06722 | 0.13279 | 0.09732 |
| RT hippocampus-PFC46 12mo | 0.19853 | 0.10712 | 0.74547 | 0.07679 |
| LT hippocampus-PFC46 12mo | 0.05102 | -0.11366 | 0.150869 | 0.07209 |
| RT TEO-Putamen 12mo | 0.19158 | 0.10204 | -0.02267 | 0.07764 |
| LT TEO-Putamen 12mo | 0.05161 | 0.10220 | 0.02511 | 0.07946 |
| RT TEO-Caudate 12mo | 0.05188 | -0.03175 | 0.18854 | 0.16421 |
| LT TEO-Caudate 12mo | 0.00804 | 0.01805 | 0.00834 | 0.12994 |
| RT V1-Putamen 12mo | 0.05230 | 0.07868 | 0.10253 | -0.07566 |
| LT V1-Putamen 12mo | 0.11521 | 0.17489 | 0.07734 | -0.11611 |

**PFC = Prefrontal cortex; OFC = Orbital frontal cortex; V = visual area; RT = Right hemisphere; LT = Left hemisphere; 12mo = 12 months of age**
