## Supplemental Table 2 for "Long-term alterations in brain and behavior after postnatal Zika virus infections in infant macaques" (from another version of this preprint: version 1, 2019-10-01)

**Supplemental Table 2.**
**Relative Volume of Brain Structures Corrected by Total Brain Volume (TBV)**

| | Subject | | | |
|---|---|---|---|---|
| **Brain Area** | **Control-1** | **Control-2** | **ZIKV-5** | **ZIKV-6** |
| TBV 12mo ($mm^3$) | 73399.37 | 84333.91 | 90258.40 | 74639.68 |
| Total CSF12mo ($mm^3$) | 11001.10 | 11709.40 | 11919.00 | 10745.40 |
| Total Intracranial Volume ($mm^3$) | 84400.47 | 96043.31 | 102177.40 | 85005.08 |
| Total Lateral Ventricle 12mo ($mm^3$) | 212.53 | 256.25 | 340.50 | 443.24 |
| Total White Matter (WM)/TBV 12mo | 29.8880 | 30.0465 | 29.6811 | 28.8612 |
| Total Grey Matter (GM)/TBV 12mo | 68.3861 | 68.2942 | 68.5905 | 69.4095 |
| Total corpus callosum/TBV 12mo | 1.0800 | 1.0285 | 1.1019 | 0.8875 |
| RT cingulate/TBV 12mo | 1.4214 | 1.4488 | 1.4252 | 1.2734 |
| LT cingulate/TBV 12mo | 1.4297 | 1.5371 | 1.4481 | 1.3536 |
| Total Amygdala/TBV 12mo | 0.7120 | 0.6441 | 0.4623 | 0.5321 |
| Total Hippocampal/TBV 12mo | 1.3488 | 1.2570 | 1.0513 | 1.1486 |
| Total Caudate/TBV 12mo | 1.6539 | 1.4213 | 1.3879 | 1.6124 |
| Total Putamen/TBV 12mo | 2.6849 | 2.5851 | 2.2376 | 2.2612 |
| Total Cerebellum GM/TBV 12mo | 5.9876 | 6.4371 | 5.3688 | 5.5499 |
| Total Cerebellum WM/TBV 12mo | 3.3330 | 3.3088 | 2.8897 | 3.4664 |
| RT prefrontal GM/TBV 12mo | 3.0989 | 2.7317 | 3.2436 | 3.6046 |
| LT prefrontal GM/TBV 12mo | 3.3523 | 3.1865 | 3.4351 | 3.6961 |
| RT prefrontal WM/TBV 12mo | 0.7397 | 0.5997 | 0.7868 | 0.9645 |
| LT prefrontal WM/TBV 12mo | 0.7969 | 0.8090 | 0.9260 | 0.9997 |
| RT frontal GM/TBV 12mo | 4.4861 | 4.6806 | 4.4204 | 4.3097 |
| LT frontal GM/TBV 12mo | 4.4990 | 4.4473 | 4.3250 | 4.4137 |
| RT frontal WM/TBV 12mo | 1.7068 | 1.8853 | 1.8290 | 1.6688 |
| LT frontal WM/TBV 12mo | 1.6457 | 1.7343 | 1.7029 | 1.5992 |
| RT parietal GM/TBV 12mo | 4.6955 | 5.2233 | 5.0329 | 4.8018 |
| LT parietal GM/TBV 12mo | 4.6311 | 4.8002 | 4.9160 | 4.6212 |

| | | | | |
|---|---|---|---|---|
| RT parietal WM/TBV 12mo | 1.8951 | 2.0800 | 2.0286 | 1.8083 |
| LT parietal WM/TBV 12mo | 1.9190 | 1.9844 | 1.9667 | 1.7048 |
| RT occipital GM/TBV 12mo | 5.55897 | 5.1850 | 6.0033 | 6.0242 |
| LT occipital GM/TBV 12mo | 5.6074 | 4.7488 | 5.6875 | 6.1439 |
| RT occipital WM/TBV 12mo | 2.3575 | 2.5353 | 2.2570 | 2.0969 |
| LT occipital WM/TBV 12mo | 2.2218 | 2.2068 | 2.0854 | 1.8585 |
| RT temporal auditory GM/TBV 12mo | 2.6791 | 2.6694 | 2.6412 | 2.6922 |
| LT temporal auditory GM/TBV 12mo | 2.6122 | 2.8163 | 2.7998 | 2.9171 |
| RT temporal auditory WM/TBV 12mo | 0.4833 | 0.5057 | 0.4976 | 0.4796 |
| LT temporal auditory WM/TBV 12mo | 0.5171 | 0.5050 | 0.5606 | 0.5216 |
| RT temporal visual GM/TBV 12mo | 3.6950 | 3.5612 | 3.6545 | 4.1533 |
| LT temporal visual GM/TBV 12mo | 4.2582 | 3.9641 | 4.1766 | 4.3187 |
| RT temporal visual WM/TBV 12mo | 1.5166 | 1.4595 | 1.4967 | 1.5667 |
| LT temporal visual WM/TBV 12mo | 1.5635 | 1.3299 | 1.5579 | 1.5066 |
| RT temporal limbic GM/TBV 12mo | 1.7058 | 1.5913 | 1.5107 | 1.3859 |
| LT temporal limbic GM/TBV 12mo | 1.4786 | 1.4933 | 1.4188 | 1.2946 |
| RT temporal limbic WM/TBV 12mo | 0.2828 | 0.2940 | 0.2773 | 0.2930 |
| LT temporal limbic WM/TBV 12mo | 0.2537 | 0.3032 | 0.2549 | 0.2601 |

**TBV = Total Brain Volume; WM = White Matter; GM = Grey Matter; CSF = Cerebrospinal fluid; RT = Right hemisphere; LT = Left hemisphere; 12mo = 12 months of age. The correction used was specific brain region/TBV x 100.**
